# BNT162b2 mRNA COVID-19 vaccine induces antibodies of broader cross-reactivity than natural infection but recognition of mutant viruses is up to 10-fold reduced

**DOI:** 10.1101/2021.03.13.435222

**Authors:** Xinyue Chang, Gilles Sousa Augusto, Xuelan Liu, Thomas M Kündig, Monique Vogel, Mona O. Mohsen, Martin F. Bachmann

**Author notes:** Correspondence: Martin Bachmann, University Clinic of Rheumatology and Immunology, University Hospital Bern, Salihaus 2, CH-3010 Bern, Switzerland; Mona Mohsen, University Clinic of Rheumatology and Immunology, University Hospital Bern, Salihaus 2, CH-3010 Bern, Switzerland. equal contribution (Mona O. Mohsen, Martin F. Bachmann).

## Abstract

**Background:** Several new variants of SARS-CoV-2 have emerged since fall 2020 which have multiple mutations in the receptor binding domain (RBD) of the spike protein.

**Objective:** We aimed to assess how mutations in RBD affected recognition of immune sera by antibodies induced by natural infection versus immunization with BNT162b2, a mRNA-based vaccine against COVID-19.

**Methods:** We produced SARS-CoV-2 RBD mutants with single mutations in the receptor binding domain (RBD) region (E484K, K417N, N501Y) or with all 3 mutations combined, as occurring in the newly emerged variants B.1.351 (South Africa) and P.1 (Brazil). Using standard and avidity ELISAs, we determined the binding capacities to mutant RBDs of antibodies induced by infection versus vaccination.

**Results:** These binding assays showed that vaccination induced antibodies recognize both wildtype and mutant RBDs with higher avidities than those raised by infection. Nevertheless, recognition of mutants RBD_K417N_ and RBD_N501Y_ was 2.5-3-fold reduced while RBD_E484K_ and the triple mutant were 10-fold less well recognized, demonstrating that the mutation at position 484 was key for the observed loss in cross-reactivity.

**Conclusion:** Our binding data demonstrate improved recognition of mutant viruses by BNT162b2-induced antibodies compared to those induced by natural infection. Recognition may, however, be 10-fold reduced for the variants B.1.351/P.1, suggesting that the development of a new vaccine is warranted. The E484K mutation is an key hurdle for immune recognition, convalescent plasma and monoclonal antibody therapy as well as serological assays based on the wildtype sequence may therefore seriously impaired.

**Capsule summary:** BNT162b2 mRNA COVID-19 vaccine-induced antibodies recognize mutant viruses with up to 10-fold lower efficiency

## Main Text

### Introduction

Because of its critical role in virus attachment and cell entry, the receptor biding domain (RBD) of SARS-CoV-2 spike (S) glycoprotein is the primary target for neutralizing antibodies protective against COVID-19^1, 2^. In this context, it has been shown that ELISA titers against RBD correlate tightly with virus neutralization^3^. It has also been demonstrated that antibody responses induced by immunization against full-length S or parts of it may result in more potent and longer-lasting responses than those elicited by natural infection with the virus^4^. These data corroborate the consensus that global vaccination programs are the most promising strategy for controlling the present COVID-19 pandemic.

A worrying factor, however, has been the emergence of variants with the ability to escape immunity produced by vaccination, potentially frustrating the unprecedented efforts to quickly develop and deploy an effective vaccine against this virus. Of the thousands of mutants identified in all continents, three recently sequenced variants have attracted the attention of the scientific community due their abnormally high rates of propagation: B.1.1.7 (K417N, E484K, N501Y, D614G), emerged in the UK; B.1.351 (K417N, E484K, N501Y, D614G), from South Africa; and P.1 (K417N/T, E484K, N501Y, D614G), appeared in Brazil. Limited knowledge of presence of cross-neutralizing antibodies induced by natural infection or vaccination is a key gap in our current understanding of the spread of SARS-CoV-2^6^. It is imperative to determine the impact of these mutations on the responses induced by currently marketed vaccines.

A previous study showed that serum neutralization is not compromised by N501Y (also found in the UK strain B.1.1.7)^7^. In contrast, E484K (found in the South Africa 501Y.V2 and in the Brazilian P.1 strains) was associated with reduced neutralization^8, 9^. Interestingly, studies applying *in vitro* pressure produced similar mutations as those that occurred naturally^10^. Whether reduced neutralization was due to impaired binding was, however, not analyzed. Here we assessed the presence of such cross-reactive antibodies in convalescent sera and sera from individuals immunized with mRNA-based BNT162b2 vaccine.

## Results and Discussion

In order to analyze cross-reactive binding of antibodies induced by infection or vaccination, we generated four mutant RBDs (RBD mutant K417N (RBD_417_) RBD mutant E484K (RBD_484_), RBD mutant N501Y (RBD_501_) and an RBD version mutated at all three sites (RBD_trip_) (Fig 1A). Please note that 2 mutations, E484K, N501Y, are localized within the receptor binding motif (RBM)^11^, directly interacting with ACE2. The domains were expressed eukaryotically in 293 cells followed by purification via HIS-tag. To assess the ability of antibodies induced by vaccination or infection to bind the different mutant RBDs, ELISAs were performed using sera from 11 convalescent patients or 6 individuals immunized with BNT162b2 (day 0 and day 21; blood was taken 2 weeks after the second dose). Our data show that binding of convalescent sera was strongly reduced for RBD_417_ and RBD_501_ and essentially abolished for RBD_484_ and RBD_trip_ (Fig 1B, left panel). In contrast, BNT162b2 induced antibodies exhibited only weakly reduced binding to RBD_417_ and RBD_501_ (2.5-3-fold), but binding was 10-fold reduced to RBD_484_ and RBD_trip_ (Fig 1B, right panel). Figure 1C summarizes the antibody titers and Figure 1D quantifies the reduced binding of vaccine induced sera to the mutant RBDs compared to wild type RBD. It is interesting to note that the single mutation E484K was equally potent at reducing antibody binding as all three mutants together in RBD_trip_, indicating that the mutation E484K is particularly problematic, perhaps because it involves a change from positive to negative charge.

**Fig 1.**
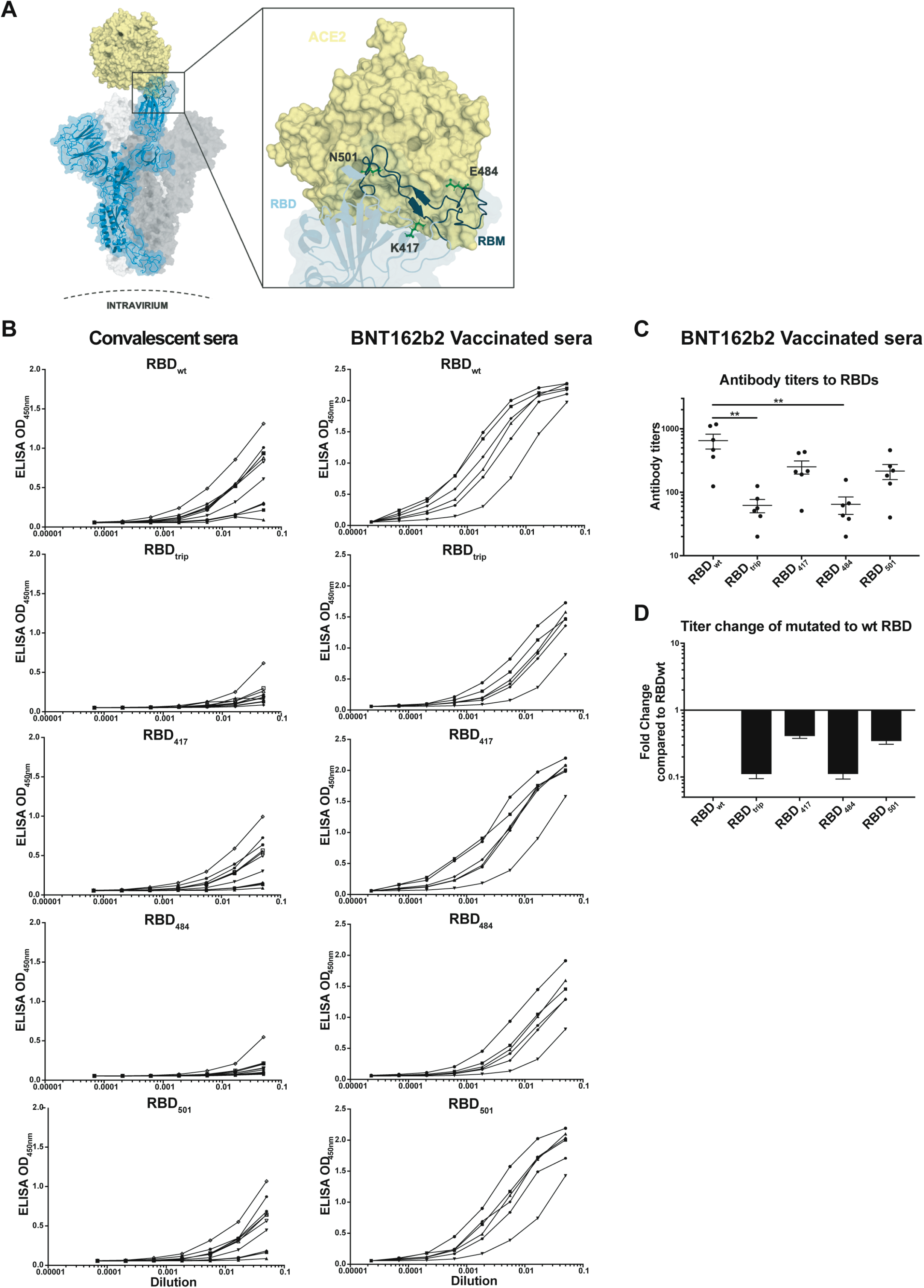
Strongly reduced recognition of mutant RBDs by convalescent and BNT162b2 vaccinated human sera. (A) Structure of RBD and location of the individual mutations used in this study (E484K, K417N, N501Y) or all 3 mutations combined). (B) Serum titration on RBDs of sera from convalescent patients (left panel) or from BNT162b2 vaccinated individuals (right panel). (C) antibody titers (OD50) of sera from 6 BNT162b2 vaccinated individuals on RBDs. (D) Fold reduction of mutant RBDs compared to wild type RBD-recognition by sera of BNT162b2 vaccinated individuals. p≤0.05 (*), p≤0.01 (**), p≤0.005 (***), p≤0.001 (****).

In order to estimate the avidity of the 2 types of antibodies for RBD and the mutants, we performed avidity assays. To this end, ELISAs were performed with an additional washing step with 7M urea and OD_50_values were compared to plates washed with PBS only. Interestingly, virus-induced antibodies were of limited avidity for wild-type RBD, and binding to mutant RBDs was essentially abolished if plates were washed with 7M urea, indicating that the antibodies that bound to the mutant RBDs were all of low avidity (Fig 2A shows results for RBD_wt_). In contrast, BNT162b2 induced antibodies were of significantly higher avidity than infection induced antibodies (Fig 2B). In addition, there was some residual binding to the mutant RBDs, indicating, however, overall low avidity as well (see below). The avidity index allows to quantify the loss in binding induced by the 7M urea wash and therefore reflect the “quality” of the antibodies. Indeed, the avidity of vaccine-induced antibodies is much higher for RBD and the mutants than those induced by infection (Fig 2C). This reduced affinity of antibodies induced by infection is consistent with the notion that individual RBDs are spaced by 25 nm on SARS-CoV-2, too large a distance for induction of optimal antibodies^12^.

**Fig 2.**
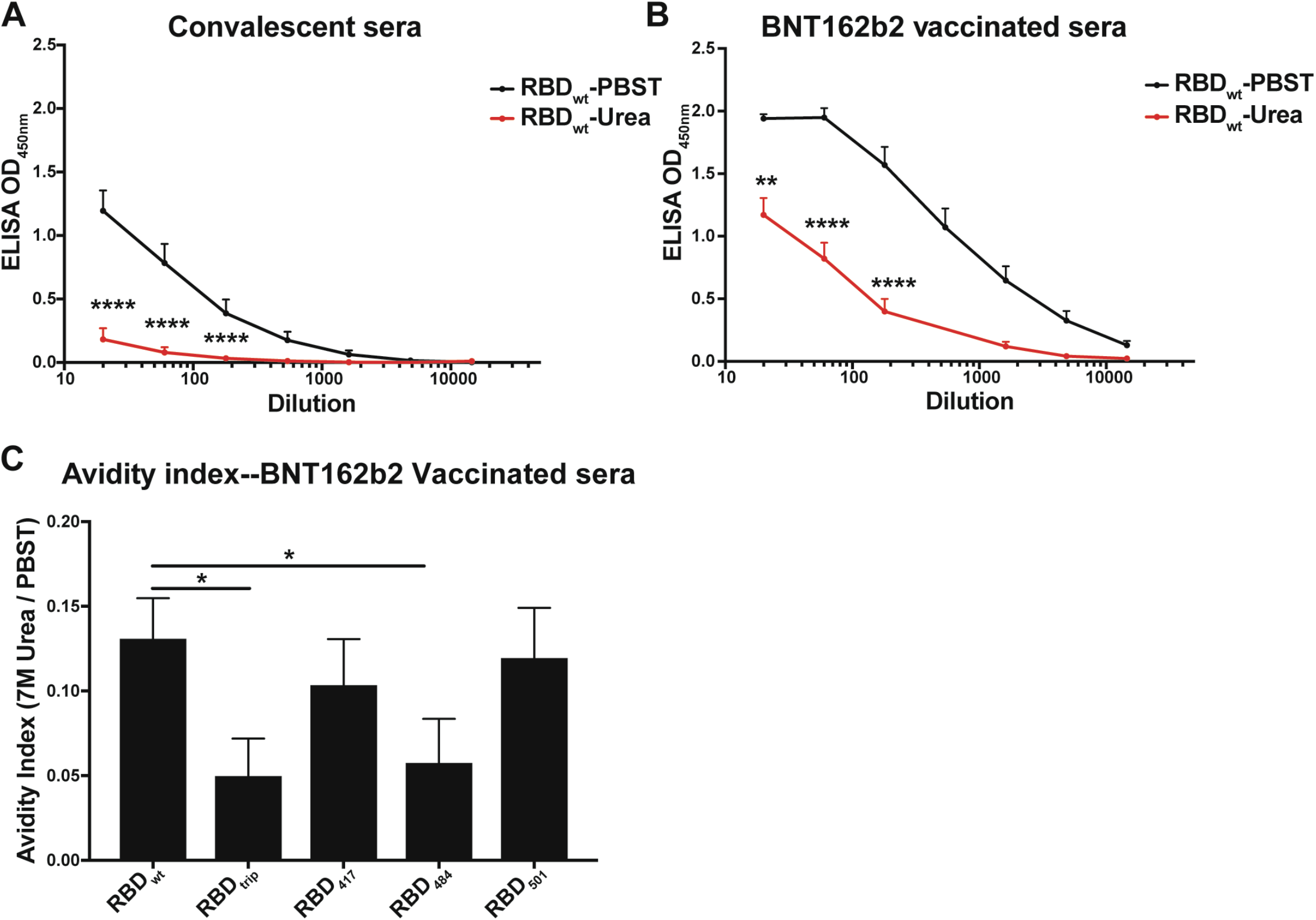
Increased avidity of BNT162b2 induced antibodies compared to antibodies induced by infection. (A,B) In order to estimate the avidity (“quality”) of induced antibodies, ELISA experiments were performed in parallel, with plates washed with PBS or 7M urea to remove low affinity antibodies (A,B) shows average recognition of wild type RBD by convalescent (A) or vaccine-induced sera (B). In black, PBS washed samples are shown, in red those washed with 7M urea. (C) Avidity indices are shown for recognition of mutant RBDs by vaccine induced sera. These indices are calculated by dividing the area under the curve of urea washed samples by the are under the curve by the PBS washed samples. Note that recognition of mutant RBDs by convalescent sera was too low to calculate a meaningful avidity index. p≤0.05 (*), p≤0.01 (**), p≤0.005 (***), p≤0.001 (****).

A key question for serology will be to see whether absence of cross-reactivity of convalescent sera holds true for individuals infected by mutant strains at position 484. This would need revision of all RBD-based serological assays.

## Abbreviations

SARS-CoV-2: Severe Acute Respiratory Syndrome Coronavirus type 2
RBD: Receptor Binding Domain
ACE2: Angiotensin-Converting Enzyme 2
RBD: Receptor Binding Domain
RBM: Receptor Binding Motif
COVID-19: Coronavirus disease 2019

## Acknowledgements

We thank Marianne Zwicker for production of mutant RBDs.

## Methods

### Protein expression and purification

The SARS-CoV-2 receptor-binding domain (RBD_wt_) and the RBD mutant (RBD_417_, RBD_501_, RBD_484_ and RBD_trip_) were expressed using Expi293F cells (Invitrogen, ThermoFisher Scientific, MA, USA). The genes that encode SARS-CoV-2 RBD_wt_ (residues Arg319-Phe541) or RBD mutants with a C-terminal 6-His-tag was inserted into pTwist CMV BetaGlobin WPRE Neo vector (Twist Bioscience, San Francisco, USA). The construct plasmids were amplified in *E*.*coli* XL-1 Blue electrocompetent cells, and were then transfected into Expi293F cells at a density of 3×10^6^ cells/ml using ExpiFectatmine 293 transfection kit (Gibco, ThermoFisher Scientific, MA, USA). The supernatant of cell culture containing the secreted RBDs was harvested 96 h after infection, and purified by passing His-Trap HP column (GE Healthcare, USA). Collected RBDs proteins were equilibrated in PBS and kept at -20°C.

### Human sera

Human sera were obtained from 11 COVID-19 convalescent patients and from 6 individuals immunized twice (days 0 and 21) with BNT162b2 according to standard protocols^13^. Sera of immunized individuals were taken 14 days after second vaccine injection.

### ELISA Assay

Corning half area 96-well plates were coated with 1 μg/ml RBD_wt_ or mutated RBDs in PBS overnight at 4°C. Then plates were blocked for 2 hours at room temperature with PBS-0.15% Casein. Afterwards, convalescent human sera, BNT162b2 immunized human sera were added, serially diluted 1:3 and incubated on plates for one hour at room temperature. Bound IgG antibodies were detected with goat anti-human IgG-POX antibody (Nordic MUbio, Susteren, The Netherlands). ELISA was developed with tetramethylbenzidine (TMB), stopped by adding equal 1 M H_2_SO_4_ solution, and read at OD_450nm_.

### Avidity ELISA

To determine the avidity of IgG antibodies, two sets of plates were prepared. Both are coated with 1 µg/ml RBDs, and after incubation with serial serum dilutions, one set of plates was washed three times for 3 minutes with 50 µl/well 7M Urea in PBS+0.05% Tween 20 whereas the other set was washed with the same amount of PBS+0.05% Tween. In-between the washing steps, all the plates were washed with PBS+0.01% Tween 20 (PBST) with the ELISA washer (BioTek, Sursee, Switzerland). The rest of the procedure is identical as described above. Avidity indexes were calculated as the fraction of the area under the curve washed with PBST versus those washed with urea.

### Data and statistical analysis

All statistical tests were performed using GraphPad PRISM 6.0 (GraphPad Software, Inc.). ELISA data in graphs are displayed as OD_450_values of individual mice ± SEM. Comparison between RBD_wt_and mutated RBDs were analyzed by paired two-tailed Student’s t-test. α=0.05 and statistical significance are displayed as p≤0.05 (*), p≤0.01 (**), p≤0.005 (***), p≤0.001 (****).

